# H3K27me3 and the PRC1-H2AK119ub pathway cooperatively maintain heterochromatin and transcriptional silencing after the loss of H3K9 methylation

**DOI:** 10.1101/2025.01.17.633676

**Authors:** Kei Fukuda, Chikako Shimura, Yoichi Shinkai

**Affiliations:** Cellular Memory Laboratory, RIKEN Cluster for Pioneering Research, Wako, 351-0198, Japan; Faculty of Life and Environmental Sciences, University of Yamanashi, Yamanashi 400-8510, Japan. Electronic address

## Abstract

Heterochromatin is a key architectural feature of eukaryotic chromosomes, essential for cell type-specific gene expression and genome stability. In the mammalian nucleus, heterochromatin is segregated from transcriptionally active euchromatic regions (A compartments), forming large, condensed, and inactive nuclear compartments (B compartments). However, the mechanisms underlying its spatial organization remain incompletely understood. Histone H3 lysine 9 and lysine 27 trimethylation (H3K9me3 and H3K27me3) are two major epigenetic modifications that enrich constitutive and facultative heterochromatin, respectively. Previously, we found that the redistribution of H3K27me3 following the loss of H3K9 methylation contributes to heterochromatin maintenance, while the simultaneous loss of both H3K27me3 and H3K9 methylation induces heterochromatin decondensation in mouse embryonic fibroblasts. However, nearly all B compartments were preserved despite the loss of these repressive chromatin modifications. These findings suggest that other factors are responsible for maintaining B compartments under these conditions. In this study, we explored the role of another repressive chromatin modification, PRC1-mediated H2A K119 monoubiquitylation (H2AK119ub/uH2A), in maintaining heterochromatin structure following the loss of H3K9/K27 methylation. We found that uH2A and H3K27me3 independently accumulate in the B compartments after the loss of H3K9 methylation in iMEFs and cooperatively maintain heterochromatin. Our data indicates that the PRC1-uH2A pathway contributes to maintain heterochromatin organization following the loss of H3K9/K27 methylation in mammalian cells.

## Introduction

The eukaryotic genome is divided into two domains: euchromatin and heterochromatin. Heterochromatin appears as an electron-dense structure in an electron microscopic image of the nucleus, which is distinct from the looser form of euchromatin and plays essential roles in genome stability, chromosome segregation, and silencing of cell type-specific gene expression (Allshire & Madhani, 2018). Heterochromatin is spatially segregated from euchromatin and is frequently found in the nuclear periphery. Recent developments in chromosome conformation capture techniques such as Hi-C have revealed that euchromatin and heterochromatin form active (A) and inactive (B) compartments, respectively (Lieberman-Aiden et al., 2009). Although the A/B compartments correlate well with gene activation/repression, the molecular determinants of compartmental forces remain largely unknown.

The epigenetic hallmark of heterochromatin, particularly constitutive heterochromatin, is the trimethylation of histone H3 lysine 9 (H3K9me3). Heterochromatin protein 1 (HP1) is a reader molecule for H3K9 methylation; it condenses chromatin (Azzaz et al., 2014; Bannister et al., 2001; Hiragami-Hamada et al., 2016; Lachner, O’Carroll, Rea, Mechtler, & Jenuwein, 2001; Zeng, Ball, & Yokomori, 2010) and drives B compartment formation in *Drosophila* embryos (Zenk et al., 2021). In mammals, at least five enzymes–SUV39H1, SUV39H2, SETDB1, G9a, and GLP– contribute to the catalysis of H3K9 methylation. SUV39H1, SUV39H2, and SETDB1 mainly catalyse H3K9me3 formation, whereas H3K9me2 formation is regulated by all five enzymes (Fukuda et al., 2021; Montavon et al., 2021; Padeken, Methot, & Gasser, 2022). Recently, we established *Setdb1/Suv39h1/Suv39h2/Ehmt1/Ehmt2* KO immortalized mouse embryonic fibroblasts (5KO iMEFs) which completely lost H3K9 methylation (Fukuda et al., 2023b). In 5KO iMEFs, H3K27me3, mediated by EZH1/2, components of the polycomb repressive complex 2 (PRC2), is redistributed to B compartments where H3K9 methylation originally accumulates. The H3K27 methyltransferase EZH1/2 dual enzymatic activities inhibitor DS3201 (DS) treatment to 5KO iMEFs (5KO+DS iMEFs) leads to weakened A/B compartmentalization, loss of electron-dense chromatin, reduced interaction with the nuclear lamina (Fukuda et al., 2023a). This indicates that H3K27me3 functions as an alternative to maintain heterochromatin in the absence of H3K9 methylation. Regardless of significant weakened A/B compartmentalization in 5KO+DS, the B compartments are still largely maintained (Fukuda et al., 2023a). We also found that H2A K119 monoubiquitylation (H2AK119ub/uH2A), which mediated by Ring1a/b, components of PRC1, also redistributes to the B compartments in 5KO iMEFs and is maintained after the depletion of H3K27me3 (Fukuda et al., 2023a). As cPRC1 and PRC1-mediated uH2A compact chromatin structure (Kim & Kingston, 2020; Zhao et al., 2024), uH2A may maintain heterochromatin after the loss of both H3K9 methylation and H3K27me3. In this study, we induced depletion of both H3K27me3 and uH2A in 5KO iMEFs and analysed transcription and 3D genome to investigate the role of repressive chromatin modifications in such cell lines. This study is the first to analyse cells that simultaneously lack major repressive chromatin modifications, including H3K9 methylation, H3K27me3, and uH2A. Through this research, we were able to comprehensively evaluate the critical roles these repressive chromatin modifications play in heterochromatin formation.

## Results

### Independent gene repression by H3K27me3 and uH2A following the loss of H3K9 methylation

We previously established *Setdb1, Suv39h1, Suv39h2, Ehmt1,* and *Ehmt2 five*-KO (5KO) iMEFs and 5KO plus *Ring1b* KO, *six*-KO (6KO) iMEFs (Supplementary Fig. S1A, Fukuda et al., 2023a). In this study, we achieved co-depletion of H3K9me, H3K27me3 and uH2A by treatment of both shRNA for *Ring1a* and DS3201 (hereafter referred to as DS) to 6KO iMEFs (Supplementary Fig. S1B-D). To investigate the function of H3K27me3 and uH2A in heterochromatin maintenance after the loss of H3K9 methylation, we performed H3K9me3/H3K27me3/uH2A ChIP-seq, RNA-seq, and Hi-C analysis. First, we analysed RNA-seq data to investigate the impact of H3K9me3/H3K27me3/uH2A depletion on the transcriptome. The principal component analysis (PCA) of gene expression levels showed that further depletion of uH2A in 5KO+DS iMEFs induced additional changes in the transcriptome, revealing that the uH2A pathway maintains the transcriptome independently of H3K27me3 after the loss of H3K9 methylation (Fig. 1A). However, the number of differentially expressed genes (DE genes: adj. P-value < 0.05, log2(FC) > 2) was larger in 5KO+DS iMEFs compared to 6KO+sh1A+DS iMEFs (Supplementary Fig. S2A). This inconsistency might be due to the use of two clones in 6KO iMEFs, which could introduce clonal variation (Fig. 1A) and make it more difficult to identify DE genes.

**Figure 1.**
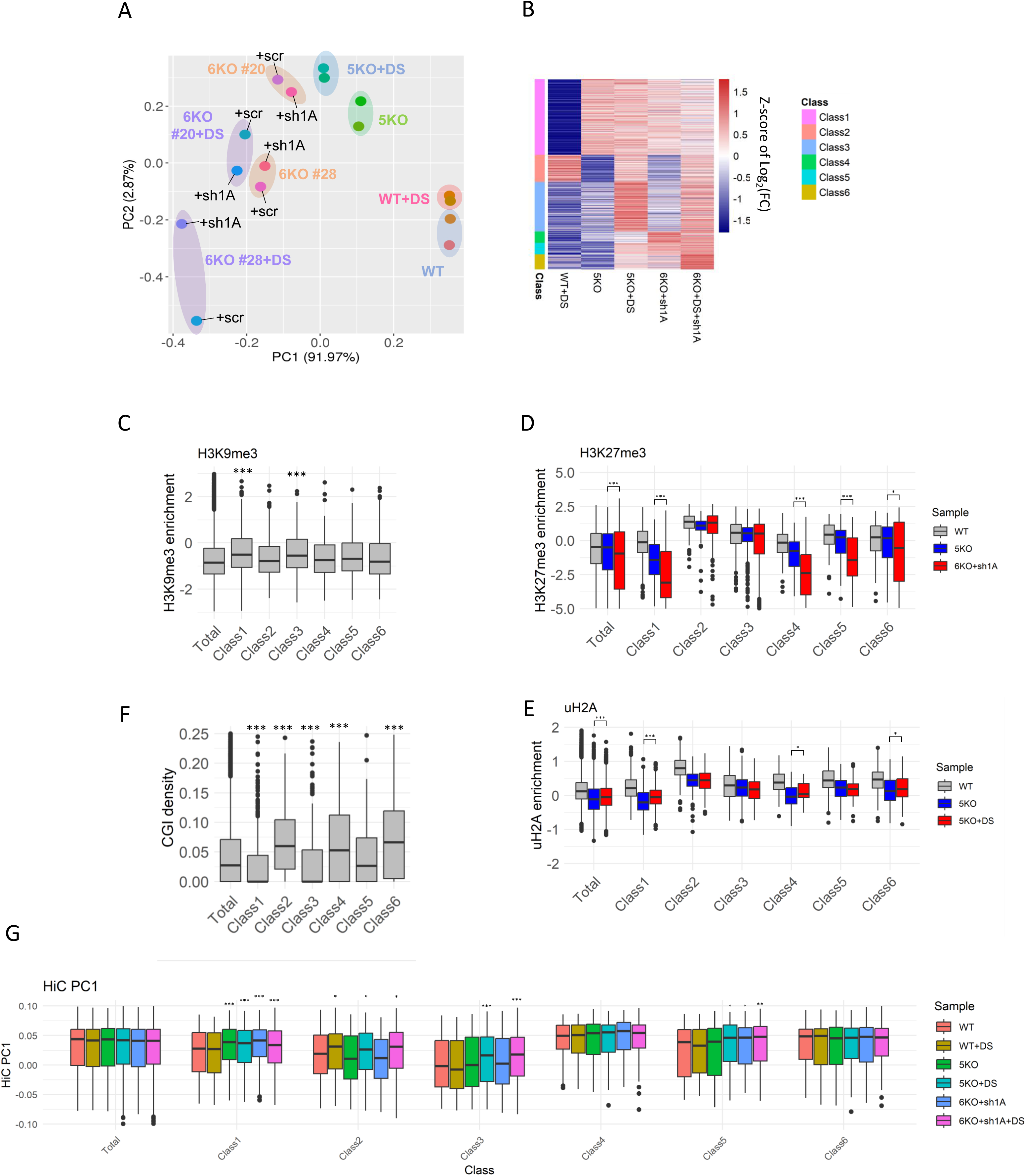
Regulation of gene repression by the H3K27me3 and RING1A/B after the loss of H3K9 methylation. (A) PCA plot of gene expression data. The PCA analysis plot based on log2(RPM) values shows the greatest expression changes in 6KO + sh1A + DS iMEF. (B) Heatmap showing expression changes across clusters. (C-E) Boxplots of histone modification enrichment. Boxplots illustrate the enrichment (Log2(ChIP/Input)) of H3K9me3 (C), H3K27me3 (D) and uH2A (E) enrichment within 10-kb around the TSS of upregulated genes in each cluster. (F) Boxplot of CGI density. This boxplot shows CGI density within 10-kb around the TSS of upregulated genes in each cluster. (G) Boxplot of Hi-C PC1 values. The boxplot depicts Hi-C PC1 values of upregulated genes in each cluster in WT iMEFs. The significance of the increase in Hi-C PC1 in the upregulated gene set was tested using a P-value. P-values in Figure 1 were calculated using Tukey’s test. Statistical significance is indicated as *, **, and *** for thresholds of 0.05, 0.01, and 0.001, respectively.

To investigate which chromatin modifications regulate the DE genes, we classified the DE genes into the following five categories: Class 1 genes: differentially expressed in 5KO iMEFs, but not in WT+DS iMEFs, Class2 genes: differentially expressed in WT+DS iMEFs, but not in 5KO iMEFs, Class 3 genes: differentially expressed in 5KO+DS iMEFs, but not in WT+DS iMEFs, 5KO and 6KO+sh1A iMEFs (regulated only by H3K27me3 in 5KO iMEFs), Class 4 genes: differentially expressed in 6KO+sh1A iMEFs, but not in WT+DS, 5KO and 5KO+DS iMEFs (regulated only by Ring1a/b in 5KO iMEFs), Class 5 genes: differentially expressed in both 5KO+DS and 6KO+sh1A iMEFs, but not in WT+DS and 5KO iMEFs (regulated by H3K27me3 and Ring1a/b inter-dependent pathway in 5KO iMEFs), Class 6 genes: differentially expressed only in 6KO+sh1A+DS iMEFs (independently regulated by both H3K27me3 and Ring1a/b) (Fig. 1B and Supplementary Fig. S2B, C). Consistent with the fact that H3K9 methylation, H3K27me3, and uH2A are repressive chromatin modifications, the number of upregulated genes exceeded that of downregulated genes in all classes (Supplementary Fig. S2A). Therefore, we focused our subsequent analyses on the upregulated genes.

Next, we analysed H3K9me3, H3K27me3, and uH2A enrichment around the TSS of DE genes (< 5kb) to investigate the regulatory mechanisms of these genes. H3K9me3 levels in Class 1 and Class 3 genes were significantly higher than those in total genes (Fig. 1C). Unlike Class 1 genes, Class 3 genes retain higher H3K27me3 in 5KO iMEFs (Fig. 1D).

Additionally, Class 3 genes were synergistically upregulated by the loss of both H3K9 methylation and H3K27me3 (Supplementary Fig. S2C), indicating that they are independently repressed through these two pathways. While H3K9me3 was not enriched in Class 4 to Class 6, a slight increase in gene expression was already observed in 5KO iMEFs, with a further increase seen in 5KO+DS compared to WT+DS or 5KO iMEFs (Supplementary Fig. S2C). This observation suggests that H3K9 methyltransferases regulate these gene groups either directly or indirectly.

The dynamics of H3K27me3 and uH2A state in depletion of H3K9 methylation or H3K9 methylation +H3K27me3 and/or uH2A were varied significantly among classes. Class 4 genes, which are suppressed by Ring1a/b, show a Ring1a/b depletion-dependent decrease in H3K27me3 levels, while uH2A levels increase upon treatment with DS3201 (Fig. 1D and E). On the other hand, in Class 5 genes, which expression are mutually suppressed by the Ring1a/b and H3K27me3 pathways, a significant decrease in H3K27me3 dependent on Ring1a/b depletion and no increase uH2A following DS3201 treatment on 5KO iMEFs were observed (Fig. 1D and E). In Class 6 genes, which transcription are independently suppressed by the Ring1a/b and H3K27me3 pathways, the decrease in H3K27me3 due to Ring1a/b depletion was less pronounced compared to Class4/5 genes, and an increase in uH2A was observed following DS3201 treatment (Fig. 1D and E). These results indicate that, in the absence of H3K9 methylation, the modes of epigenomic and transcriptional regulation by the Ring1a/b and H3K27me3 pathways vary depending on the gene.

To elucidate the molecular basis of transcriptional regulation diversity mediated by the Ring1a/b and H3K27me3 pathways in 5KO iMEFs, we investigated whether differences in genomic sequences or the spatial organization of the genome underlie these variations. The analysis showed that genes suppressed by the H3K27me3 pathway in WT iMEFs (Class 2) was characterized by high CGI density around their TSS and mainly located in the A compartments. The class suppressed solely by H3K27me3 in 5KO iMEFs (Class 3) exhibited the lowest CGI density among PRC-related upregulated classes (Class 2-6) and was preferentially located in the B compartments (Fig. 1F and G). The classes where the Ring1a/b pathway represses independently of the H3K27me3 pathway in 5KO iMEFs (Class 4/6) were characterized by high CGI density and strong A compartments (Fig. 1F and G). The classes where the H3K27me3 and Ring1a/b pathways mutually depend on each other for repression (Class 5) exhibit intermediate features between those of Class 3 and Class 4/6 (Fig. 1F and G). These results suggest that the composition of genome sequences and nuclear compartments contribute to differences in repression systems among genes. Furthermore, genes in Class 1, 2, 3, and 5 showed an increase in compartment scores correlated with elevated gene expression, whereas this correlation was not observed in Class 4 and Class 6 (Fig. 1G and Supplementary Fig. S2D). Therefore, it is suggested that the Ring1a/b pathway suppresses transcription within strong A compartments (class 4 and 6 genes) without inducing changes in compartmentalization.

### Redistribution of H3K27me3 and uH2A after the loss of H3K9 methylation contributes to transposon repression

It has been reported that the deposition of H3K27me3 on transposons contributes to their silencing in female primordial germ cells, where H3K9me3 levels are low (Huang et al., 2021). That prompted us to investigate transposon expression in our cell lines. More than 100 types of transposons showed increased expression in 5KO iMEFs, and when H3K27me3 and/or uH2A were further depleted, the expression of additional transposons was increased (Supplementary Fig. S3A). In contrast, transposon downregulation was detected solely in 5KO iMEFs, and it was limited to MER68B. This suggests that H3K9me, H3K27me3, and uH2A play a fundamental role in suppressing transposons. To clarify the suppressing mechanisms of transposons by in our cell lines, we classified them in the same way as genes with increased expression. As a result, 113, 0, 42, 14, 41 and 25 transposon types were categorized into each Class 1 to 6, respectively (Fig. 2A, Supplementary Fig. S3B). In all classes, the majority were LTR-type transposons (Fig. 2B). Since H3K9me3 accumulated in transposons across all classes, it suggests that transposons with increased expression are primarily targets of H3K9me3 pathway (Fig. 2C). Although DS3201 treatment did not induce transposon activation in WT iMEFs (Class 2), it did result in transposon activation in 5KO iMEFs (Class 3), suggesting that the H3K27 methylation pathway functions compensatory to suppress transposons in the loss of H3K9 methylation, as previously reported (Huang et al., 2021). In addition to the H3K27me3 pathway, Ring1a/b also contributed to transposon repression either in coordination with or independently of the H3K27me3 pathway (Class 4-6), revealing that transposons are regulated through a multilayered suppression system (Fig. 2A and Supplementary Fig. S3B).

**Figure 2.**
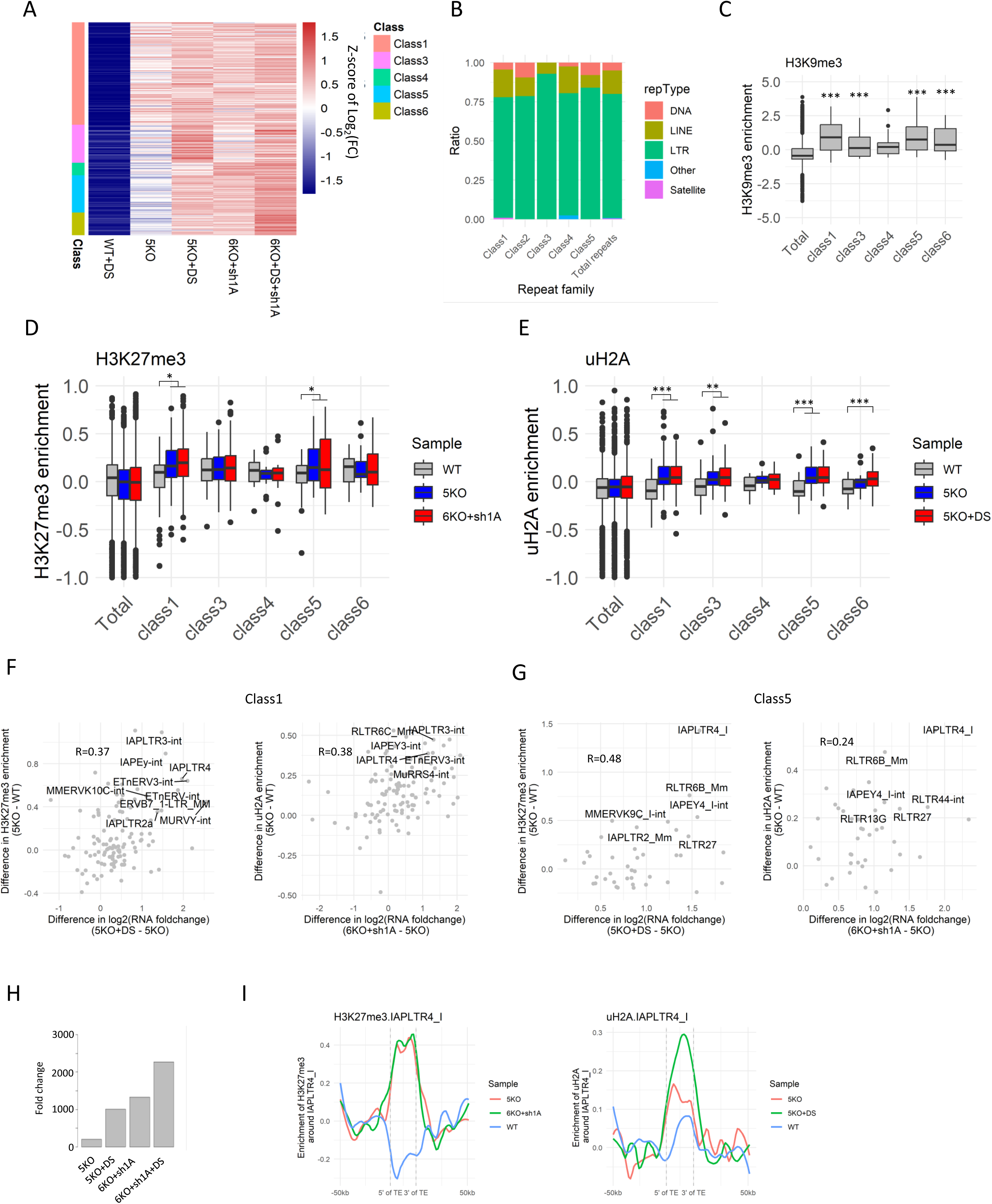
Regulation of transposon repression by the H3K27me3 and RING1A/B after the loss of H3K9 methylation. (A) Heatmap showing expression changes of upregulated transposons across clusters. (B) Fraction of repeat type in each Class of upregulated transposons. (C-E) Boxplots of histone modification enrichment. Boxplots illustrate the enrichment (Log2(ChIP/Input)) of H3K9me3 (C), H3K27me3 (D) and uH2A (E) enrichment in upregulated transposons in each cluster. (F-G) Scatter plots of epigenome changes and transcriptional changes in Class 1 (F) and class 5 (G) transposons. Left: The x-axis represents expression differences between 5KO iMEFs and 5KO+DS iMEFs. The y-axis represents H3K27me3 enrichment differences between WT iMEFs and 5KO iMEFs. Right: The x-axis represents expression differences between 5KO iMEFs and 6KO+sh1A iMEFs. The y-axis represents uH2A enrichment differences between WT iMEFs and 5KO iMEFs. Pearson’s coefficient R indicates the correlation between epigenome change and transcriptional change. (H) Bar plot represents the fold change of IAPLTR4_I in each sample. (I) Enrichment of H3K27me3 and uH2A around IAPLTR4_I in each sample. P-values in Figure 2 were calculated using Tukey’s test. Statistical significance is indicated as *, **, and *** for thresholds of 0.05, 0.01, and 0.001, respectively.

Epigenome analysis of transposons revealed an increased accumulation of H3K27me3 and uH2A on Class 1 and Class 5 transposons in 5KO iMEFs compared to WT iMEFs (Fig. 2D and 2E). This accumulation was accompanied by increased expression of these transposons in 5KO + DS and 6KO + sh1A iMEFs compared to 5KO iMEFs (Fig. 2A). Notably, the elevated expression of Class 1 and Class 5 transposons correlated with enhanced enrichment of H3K27me3 and uH2A in 5KO iMEFs (Fig. 2F and 2G). Among the Class 5 transposons, IAPLTR4_I was already upregulated in 5KO iMEFs but showed further increases in expression in 5KO + DS and 6KO + sh1A iMEFs, with the highest levels observed in 6KO + sh1A + DS iMEFs (Fig. 2H). H3K27me3 and uH2A enrichment on IAPLTR4_I was significantly elevated in 5KO iMEFs compared to WT iMEFs and was maintained independently of each other (Fig. 2I). These findings suggest that H3K27me3 and uH2A are redistributed to IAPLTR4_I following the loss of H3K9 methylation, where they independently contribute to the suppression of this transposon. To summarize the above analysis, our findings indicate that the loss of repressive chromatin modifications on transposons triggers the redistribution of other repressive chromatin marks, ensuring the redundant maintenance of the repressive state.

### H3K27me3 and uH2A redistribute to the B compartments to maintain nuclear compartments after the loss of H3K9 methylation

From the above analysis, it was demonstrated that the redistribution of H3K27me3 and uH2A following the loss of H3K9 methylation contributes to the maintenance of transcriptional repression of retroelements. To capture this redistribution phenomenon more globally, we analysed the accumulation of H3K27me3 and uH2A at 250-kb bin. In WT iMEFs, H3K27me3 and uH2A were largely correlated (R=0.54) and were mainly enriched in the A compartments (Fig. 3A). In 5KO iMEFs, the correlation between H3K27me3 and uH2A was increased (R=0.87) compared to WT iMEFs, and both H3K27me3 and uH2A were enriched in the B compartment (Fig. 3B). Therefore, following the loss of H3K9 methylation, H3K27me3 and uH2A undergo a similar redistribution and accumulate in the B compartment. Ring1a/b depletion in 5KO iMEFs did not affect H3K27me3 distribution (Fig. 3C). Similarly, uH2A distribution in 5KO iMEFs were not largely affected by DS3201 treatment to 5KO iMEFs (Fig. 3D). However, the overall impression is the redistributed H3K27me3 and uH2A in the B compartments are relatively independent of each other in this cell population. In the genomic regions where H3K27me3 was redistributed (log2(ChIP/Input) < 0 in WT and log2(ChIP/Input) > 0 in 6KO+sh1A), transcription levels and Hi-C PC1 values were increased from 6KO to 6KO+sh1A+DS (Fig. 3E and 3F). Similarly, in the genomic regions where uH2A was redistributed, transcription levels and Hi-C PC1 values were increased from 5KO+DS to 6KO+sh1A+DS (Fig. 3G and 3H). Therefore, both H3K27me3 and uH2A redistribute to the B compartment after the loss of H3K9 methylation, maintaining transcriptional repression and the B compartment status, as observed in Fig. 3I.

**Figure 3.**
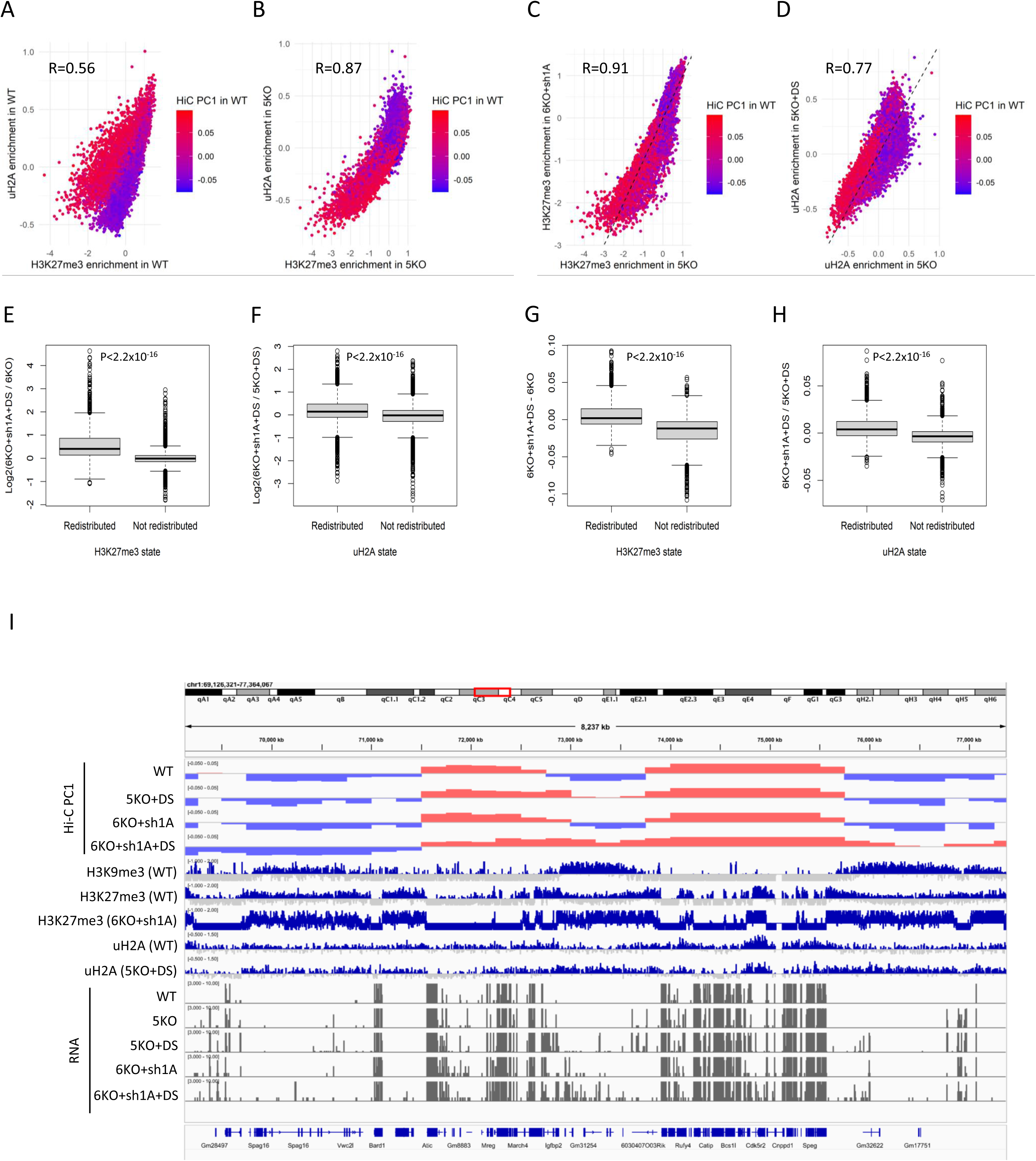
Function of redistributed H3K27me3 and uH2A in 5KO iMEFs. (A, B) Scatter plots showing the enrichment of H3K27me3 and uH2A in WT (A) and 5KO iMEFs (B). Each plot is coloured according to the Hi-C PC1 value in WT iMEFs and represents the log2(ChIP/Input) for 250-kb bins. R denotes the Pearson correlation coefficient. (C) Scatter plots showing the enrichment of H3K27me3 in 5KO iMEFs and 6KO+sh1A iMEFs. (D) Scatter plots showing the enrichment of uH2A in 5KO iMEFs and 5KO+DS iMEFs. (E) Box plot showing changes in transcription levels between 6KO+sh1A iMEFs and 6KO+sh1A+DS iMEFs. The analysis compares regions where H3K27me3 has been redistributed (log2(ChIP/Input) < 0 in WT iMEFs and > 0 in 6KO+sh1A iMEFs) to regions where redistribution has not occurred within 250-kb bins (log2(ChIP/Input) < 0 in WT iMEFs and 6KO+sh1A iMEFs). (F) Box plot showing changes in transcription levels between 5KO+DS iMEFs and 6KO+sh1A+DS iMEFs. The analysis compares regions where uH2A has been redistributed (log2(ChIP/Input) < 0 in WT iMEFs and > 0 in 5KO+DS iMEFs) to regions where redistribution has not occurred within 250-kb bins (log2(ChIP/Input) < 0 in WT iMEFs and 5KO+DS iMEFs). (G) Box plot showing changes in Hi-C PC1 values between 6KO+sh1A iMEFs and 6KO+sh1A+DS iMEFs. The analysis compares regions where H3K27me3 has been redistributed to regions where redistribution has not occurred in 250-kb bins. (H) Box plot showing changes in Hi-C PC1 values between 5KO+DS iMEFs and 6KO+sh1A+DS iMEFs. The analysis compares regions where H3K27me3 has been redistributed to regions where redistribution has not occurred in 250-kb bins. (I) Representative regions where H3K27me3 and uH2A are redistributed, maintaining transcription and nuclear compartmentalization. P-values in Figure 3 were calculated using a t-test.

### H3K27me3 and uH2A cooperatively maintain A/B compartment segregation after the loss of H3K9 methylation

Finally, we investigated global 3D genome profiles. The PCA analysis of Hi-C PC1 values showed the most significant changes in the 6KO+sh1A+DS iMEFs among samples (Supplementary Fig. S4A). The interaction matrix suggests that a marked decrease in long-range interactions in the 6KO+sh1A+DS iMEFs compared other samples (Fig. 4A), and the Pearson correlation matrix also shows weakened A/B compartment segregation in 6KO+sh1A+DS iMEFs (Fig. 4B). As expected, B-B interactions were significantly decreased in the 6KO+sh1A+DS iMEFs (Fig. 4C), and compartment strength also showed a significant decrease (Supplementary Fig. S4B). These findings indicate that H3K27me3 and Ring1a/b cooperatively maintain the segregation of A/B compartments. Unexpectedly, in 5KO iMEFs and 6KO+sh1A iMEFs, an increase in B-B interactions was observed, with the strongest interactions detected in 6KO+sh1A iMEFs. This suggests that a reduced diversity of repressive chromatin modifications facilitates tighter clustering of B compartments. Despite the significant reduction in B-B interactions in 6KO+sh1A+DS iMEFs, over 80% of the B compartment was still maintained in the condition (Supplementary Fig. S4C), it suggests that other factors are also involved in the formation of A/B compartments. Unlike the compartments, the insulation score showed no significant difference between 5KO+DS iMEFs and 6KO+DS+sh1A iMEFs, implicating that the depletion of Ring1a/b has little impact on the TAD pattern (Supplementary Fig. S4D).

**Figure 4.**
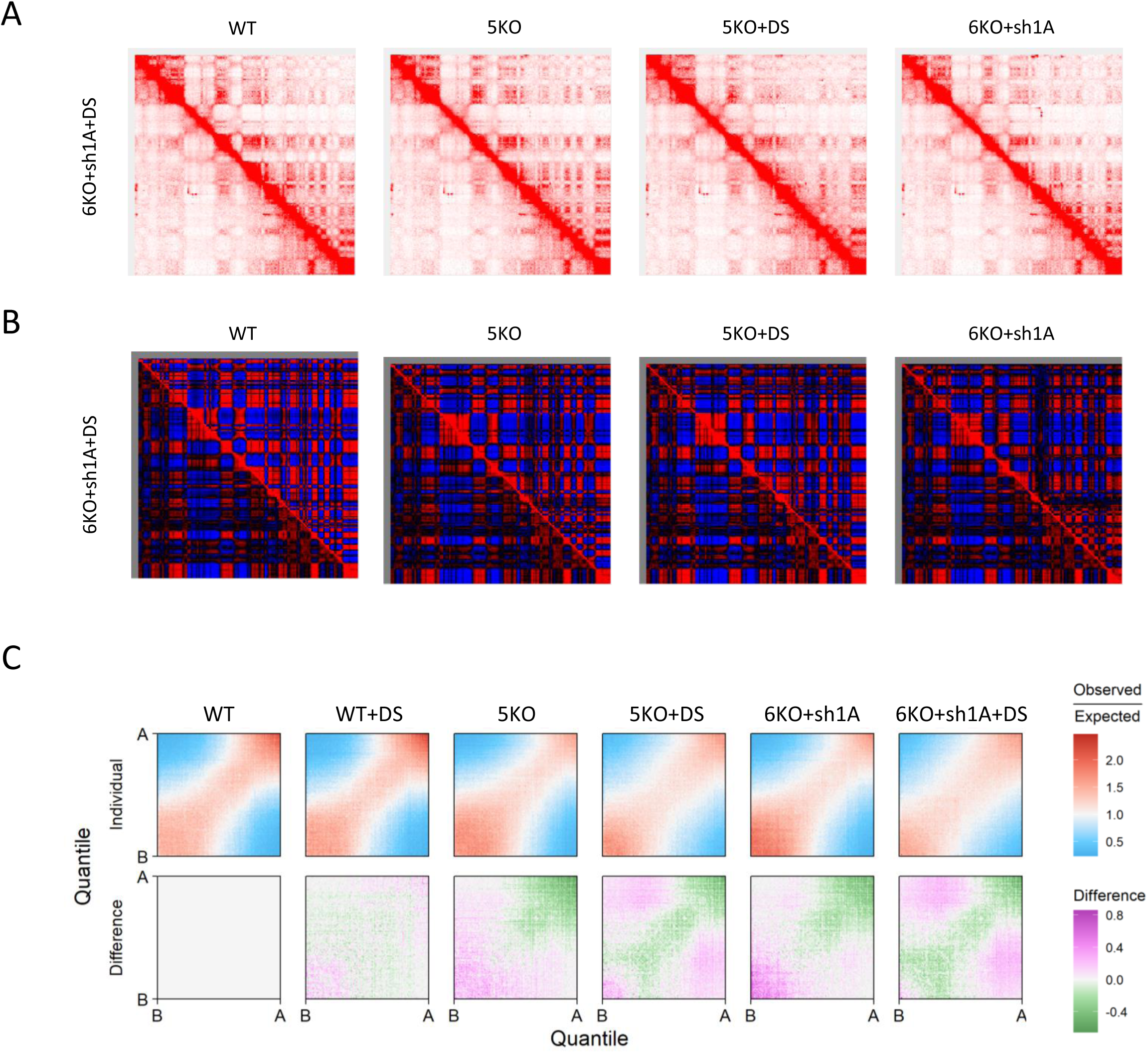
H3K27me3 and RING1A/B maintain A/B compartment segregation after the loss of H3K9 methylation. (A, B) Interaction matrix (A) and pearson correlation matrix (B) between 6KO+sh1A+DS iMEFs and other samples. (C) Saddle plots of each sample. The saddle plot shows a decreased B-to-B and A-to-A interactions and an increased A-to-B interaction in 6KO+sh1A+DS iMEFs.

## Discussion

In this study, we successfully analysed the transcription, epigenome and 3D genome organization in mammalian cells that, for the first time, lacked all three major repressive chromatin modifications: H3K9 methylation, H3K27me3, and uH2A. Our findings revealed that, following the loss of H3K9 methylation, H3K27me3 and PRC1-uH2A can independently or some inter-dependently contribute to maintain gene/TE silencing and nuclear compartments.

### After the loss of H3K9 methylation, PRC1 and PRC2 pathways regulate distinct sets of genes

From our transcriptome analysis, H3K27 methylation (H3K27me3) mediated by PRC2, suppresses genes with the high-CGI located in weak-A compartment in WT iMEFs, whereas in the absence of H3K9me, it becomes critical for suppression of low-CGI-density genes, too. Notably, in 5KO iMEFs, a group of genes (Class 3) whose transcription is suppressed solely by H3K27me3 are characterized by low CGI density. Both H3K9me3 and H3K27me3 accumulate at class 3 genes in WT iMEFs, and their expression does not increase in DS3201-treated WT iMEFs, indicating that these genes are independently repressed by both H3K9me3 and H3K27me3 in WT iMEFs. On the other hand, the group of genes repressed by PRC1 (Class 4 and 6), which mediates uH2A, lacks H3K9me3 accumulation. Although we didn’t validate the impact of PRC1 KD in WT iMEFs, these genes, characterized by high CGI and strong-A compartments, differ from those repressed by H3K9 methylation (Class 1), suggesting that H3K9me3 and PRC1 target distinct sets of genes which might be expected based on the previous analyses (Bell, Burton, Dean, Gasser, & Torres-Padilla, 2023).

The regulation of repressive chromatin modifications by PRC1 and PRC2 is closely interconnected. PRC2 assembles into two subcomplexes in mammals: PRC2.1 with polycomb-like proteins (PCLs) and PRC2.2 with JARID2 and AEBP2 (Holoch & Margueron, 2017). PRC1 complexes can be broadly subdivided into canonical and variant complexes (cPRC1 and vPRC1, respectively), based on distinct modes of genomic targeting. First, both PRC1 and PRC2 complexes possess non-methylated CpG binding components for CGI targeting, such as KDM2B for vPRC1-PCGF1 complex (Farcas et al., 2012; He et al., 2013) and PCL1/2/3 for PRC2.1 complex (Li et al., 2017), respectively. Then, cPRC1 localizes at PRC2 targets by binding to H3K27me3 via a chromobox domain–containing (CBX) protein (Cao et al., 2002; Min, Zhang, & Xu, 2003; Wang et al., 2004). PRC2.1 and PRC2.2 each recruit different types of CBX proteins within cPRC1 to their respective target regions, likely through the binding of their chromodomains to H3K27me3 (Glancy et al., 2023). vPRC1 incorporates Rybp/Yaf2 instead of CBX proteins and is recruited to target genes independently of H3K27me3 and PRC2 (Blackledge et al., 2014; Gao et al., 2012; Hauri et al., 2016; Kloet et al., 2016; Tavares et al., 2012). vPRC1 also promotes to PRC2 recruitment (Blackledge et al., 2014) might be via interaction between uH2A containing nucleosome and JARID2-AEBP2 (Kalb et al., 2014). Therefore, while PRC1 and PRC2 are closely related, their hierarchy differ depending on the complex type. vPRC1 contains KDM2B, which plays a key role in recognizing CGIs (Farcas et al., 2012; He et al., 2013). This could explain our observation that the gene set derepressed by PRC1 KD in 5KO iMEFs is enriched in CGI-rich genes. The phenomenon that PRC1 has a higher preference for CGI-containing genes than PRC2 has also been reported *in vivo* in mice. Polycomb proteins play a major role in X-chromosome inactivation (Brockdorff, 2017). In the extra-embryonic ectoplacental cones, PRC2-deficient mice caused by *Eed* KO exhibit increased expression of X-chromosomal genes lacking CGI in their promoters. In contrast, PRC1- deficient mice caused by *Ring1a/b* KO show increased expression of only CGI-containing X-linked genes (Masui et al., 2023). How PRC2 targets non-CGI genes remains largely unclear (Laugesen, Hojfeldt, & Helin, 2019), but PHF19, a Polycomb-like protein included in PRC2.1, has been reported to be crucial for the formation of H3K27me3 in non-CGI regions in the human plasma cell leukemia cell line L-363 (Ren et al., 2019). This suggests that PRC2 subtypes may play a key role in targeting non-CGI regions. The functional differences between PRC1 and PRC2, as described above, are suggested to be maintained even after the loss of H3K9 methylation, from this study.

### Repression of transposons in 5KO iMEFs through the redistribution of H3K27me3/uH2A

The gene sets repressed by the PRC1 and PRC2 pathways in 5KO iMEFs showed higher H3K27me3/uH2A level around TSS than those of total genes in WT iMEFs (Fig. 1D and E), whereas transposons exhibited redistribution of H3K27me3/uH2A after the loss of H3K9 methylation, which was found to correlate with transcriptional repression (Fig. 2D). The accumulation of H3K27me3 on transposons induced by the reduction of H3K9 methylation and DNA methylation is observed not only in mutants of these modifying enzymes but also during developmental processes, such as in mammalian primordial germ cells (Huang et al., 2021). We have demonstrated for the first time that not only H3K27me3 but also uH2A undergo redistribution to transposons after the loss of H3K9 methylation; however, it remains unclear whether the redistribution of H3K27me3 and uH2A is independent of each other. Nonetheless, since the presence of one is often maintained even when the other is lost, it is likely that they are independently maintained. The exact mechanism by which PRC1/2 recognizes transposons also remains unclear and warrants further investigation.

### Redistribution of H3K27me3/uH2A to the B compartments in 5KO iMEFs

Similarly to transposons, we have demonstrated for the first time that not only H3K27me3 but also uH2A undergo redistribution to the B compartments and contribute to the maintenance of A/B segregation. Such PRC1/2 redistribution is also observed in the pericentromeric heterochromatin (PCH) of *Suv39h1/2*-deficient cells and DNA methylation-deficient mouse ES cells (Peters et al., 2003; Saksouk et al., 2014). In *Suv39h1/2*-deficient cells, reduced DNA methylation at PCH facilitates the binding of the DNA methylation-sensitive sequence-specific DNA-binding protein BEND3, which in turn promotes the deposition of H3K27me3 at PCH (Saksouk et al., 2014).

Additionally, it has been reported that the binding of KDM2B to PCH is sufficient to recruit PRC1/2 to PCH (Cooper et al., 2014). Since DNA hypomethylation attracts PRC1/2 to CpG-rich genomic regions, the hypomethylation of PCH may promote the binding of KDM2B to PCH, leading to the recruitment of PRC1/2. In mouse zygotes, the paternal PCH exhibits reduced H3K9me3 levels, with PRC1/2 binding instead (Puschendorf et al., 2008). Importantly, even in the absence of H3K27me3, PRC1 binding to paternal PCH is maintained, suggesting that vPRC1, which contains KDM2B, rather than cPRC1, may play a critical role in recruiting PRC1 to PCH under physiological conditions. In 5KO iMEFs, DNA methylation in the B compartment is reduced (Fukuda et al., 2023b), raising the possibility that the DNA methylation-sensitive DNA-binding proteins, such as KDM2B or BEND3, mediate the redistribution of PRC1/2 to the B compartment.

### Maintenance of A/B compartment segregation by H3K27me3/uH2A in 5KO iMEFs

This study also revealed that the loss of H3K9 methylation leads to the redistribution of H3K27me3 and uH2A to the B compartment, where they independently maintain A/B compartmentalization. Polycomb group proteins have been reported to influence chromatin organization at multiple levels, including chromatin compaction, long-range interactions, and phase separation (Uckelmann & Davidovich, 2024). However, the A/B compartment separation was still largely maintained in all H3K9me/H3K27me3/uH2A- depleted cells. This could be due to several possibilities: the partial retention of uH2A caused by incomplete *Ring1a* depletion via knockdown, the potential lethality of cells if compartmentalization is entirely lost, or the involvement of other factors. For instance, the clustering of transcriptionally active regions driven by transcriptional activity may passively contribute to the formation of A/B compartments. Future studies are needed to investigate how nuclear compartments are affected by transcriptional inhibition or the disruption of active chromatin modifications.

---

**Supplementary Table S1. Summary of NGS data in this study.**

**Supplementary Table S2. List of upregulated class of genes.**

**Supplementary Table S3. List of upregulated class of transposons.**

## Materials and Methods

### Cell culture

We used previously established *Setdb1, Suv39h1/2, Ehmt1, Ehmt2, Ring1b* KO iMEFs to analysis the role of uH2A in heterochromatin maintenance (Fukuda et al., 2021). Mouse embryonic fibroblasts were maintained in Dulbecco’s modified Eagle’s medium (Nacalai tesque, 08458-16) containing 10% fetal bovine serum (Biosera, FB1061), MEM Non-Essential medium and 2-Mercaptoethanol (Nacalai tesque, 21417- 52). To inhibit EZH1/2 catalytic activity, iMEFs were cultured for seven days with 1 μM DS3201.

### *Ring1a* knockdown by shRNA

We produced a lentiviral vector expressing shRNA targeting *Ring1a* by transfecting 293FT cells with shRNA vector, psPax2, and pMD2.G using PEI. Two days later, we collected the culture supernatant and then used it to transduce 5KO-*Ring1b* KO iMEFs at MOI=2. Following selection with 7 μg/ml BSD for 3 days, we collected the cells.

### Native ChIP and crosslinked ChIP

Native ChIP assays were performed as described previously (Fukuda et al., 2021). Mouse monoclonal antibody against H3K27me3 (1E7) was used.

### H2AK119 monoubiquitylation (H2Aub) Carrier Assisted ChIP-seq (CATCH-seq)

H2Aub CATCH-seq libraries were prepared as previously described (Zhu et al., 2021) with some modifications. Briefly, 2,000 MEFs per sample were used for library construction. Cells were permeabilized by Nuclei EZ lysis buffer supplemented with 0.1% Triton, 0.1% deoxycholate, complete EDTA-free protease inhibitor cocktail and 1 mM phenylmethanesulfonyl fluoride on ice. For each sample, 800 S2 Drosophila culture cells were added for a spike-in normalization purpose. Chromatin was fragmented in situ by 2 U/µl MNase (M0247S, NEB) at 37C for 7.5 min. The reaction was stopped by adding 1/10 volume of 100 mM EDTA and diluted in the freshly prepared immunoprecipitation buffer. As an input, 5% of the chromatin lysate was taken. Then, 30 ng of annealed I-SceI carrier DNA was added to each sample. The forward and reverse strands of the carrier DNA were as follows:

/5AmMC6/Gtagggataacagggtaattagggataacagggtaattagggataacagggtaattagggataacagggtaattagggataacagggtaattagggat aacagggtaat*c/3AmMO/ and /5AmMC6/Gattaccctgttatccctaattaccctgttatccctaattaccctgttatccctaattaccctgttatccctaattaccctgttatccctaattaccctgttatcc cta*c/3AmMO/, respectively, where asterisks represent phosphorothioate bonds. The oligos were synthesized by Integrated DNA Technologies. For each immunoprecipitation reaction, 0.5 µl of rabbit monoclonal anti- H2AK119 monoubiquitylation antibody (Cell Signaling, #8240) conjugated to precleared Dynabeads Protein A and G mixture was used. After immunoprecipitation at 4C overnight, the chromatin-Dynabeads were washed by the low and high salt wash buffers, and the chromatin was eluted in the freshly prepared ChIP elution buffer at 65C for 1 hr. DNA was recovered by phenol-chloroform extraction followed by ethanol precipitation. Adaptor ligation was pre-formed by NEBNext Ultra II DNA Library Prep Kit for Illumina (E7645, NEB) in a half scale of the manufacturer’s instruction, and the libraries were purified by 1.8x SPRIselect beads (B23318, Beckman Coulter). The DNA was amplified by KAPA Hifi 2X mater PCR mix (KK2605) for 14 PCR cycles with indexing primers. After purification with 0.9x SPRIselect beads, the samples were digested by I-SceI (5 U/µl, NEB, R0694) 37C for 2 hrs followed by heat inactivation at 65C for 20 min and purified by 0.9x SPRIselect beads. The second amplification was not performed. The libraries were sequenced on a HiSeq X.

### Western blot analysis

Western blot analysis was performed as described previously (Fukuda et al., 2021). Briefly, cells were suspended in RIPA buffer (50mM Tris-HCl (pH 8.0), 420mM NaCl, 0.5% sodium deoxycholate, 0.1% Sodium dodecyl sulphate, 1% NP-40) and sonicated. The extract was incubated for 30 minutes on ice, and then incubated at 95°C for 5 minutes. The extract was loaded and run on SDS-PAGE gel as standard protocols. For histone proteins, intensity was analysed by OdysseyR CLx Imagins System(LI-COR). Anti-lamin B1 (12987-1-AP, Proteintech), anti-lamin A/C (3A6-4C11, Diagenode), anti-H2Aub (D27C4), anti-Ring1B (monoclonal antibody kindly gifted from Dr. Koseki, RIKEN), anti-Ring1A (#2820, CST) were used.

### Preparation of ChIP-seq library

The ChIP DNA was fragmented by Picoruptor (Diagenode) for 10 cycles of 30 seconds on, 30 seconds off. Then, ChIP library was constructed by KAPA Hyper Prep Kit (KAPA BIOSYSTEMS) and SeqCap Adapter Kit A (Roche) according to manufacturer instructions. The concentration of the ChIP-seq library was quantified by KAPA Library quantification kit (KAPA BIOSYSTEMS). ChIP sequencing was performed on a HiSeq X platform (Illumina). We performed two biological replicates for ChIP-seq.

### Preparation of RNA-seq library

500 ng of total RNA was used for RNA-seq library construction. RNA-seq library was constructed by KAPA mRNA Hyper Prep Kit (KAPA BIOSYSTEMS) and SeqCap Adapter Kit (Roche) according to manufacturer instructions. The concentration of the RNA-seq library was quantified by KAPA Library quantification kit (KAPA BIOSYSTEMS). mRNA sequencing was performed on a HiSeq X platform (Illumina). We performed two biological replicates for RNA-seq.

### Preparation of Hi-C library

Hi-C experiments were performed as previously described (Ikeda et al., 2018; Kadota et al., 2020), based on DpnII enzyme (4-bps cutter) using 2×10^6^ fixed cells. Hi-C libraries were subject to paired-end sequencing (150 base pair (bp) read length) using HiSeq X Ten. Detailed protocol for HiC-seq library preparation is available at Protocols.io (https://www.protocols.io/view/iconhi-c-protocol-ver-1-0-4mjgu4n). We performed two biological replicates for Hi-C.

### Hi-C data analysis

Hi-C data processing was done by using Docker for 4DN Hi-C pipeline (v43, https://github.com/4dn-dcic/docker-4dn-hic). The pipeline includes alignment (using the mouse genome, mm10) and filtering steps. After filtering valid Hi-C alignments, *.hic* format Hi-C matrix files were generated by Juicer Tools (Durand et al., 2016) using the reads with MAPQ>10. The A/B compartment (compartment score) profiles (in 250 kb bins) in each chromosome (without sex chromosome) were calculated from *.Hic* format Hi-C matrix files (intrachromosomal KR normalized Hi-C maps) by Juicer Tools (Durand et al., 2016) as previously described (Miura, Poonperm, Takahashi, & Hiratani, 2018). We averaged Hi-C PC1 values in each 250 kb bin from two biological replicates for the downstream analysis. TAD boundaries were identified as previously reported (Miura et al., 2019). Chromatin loops were identified by SIP, Significant Interaction Peak caller, v1.6.1 (Rowley et al., 2020) with the following parameters: -norm KR -g 0.5 -min 2.0 -max 2.0 -mat 2000 -d 5 -res 40000 -sat 0.01 -t 2800 -nbZero 6 -factor 1 -fdr 0.015 -del true -cpu 1 -isDroso false. We used GENOVA tools (van der Weide et al., 2021) for producing aggregate TAD plots, insulation analysis and pyramid plots. We also used Juicer Tools (Durand et al., 2016) for producing interaction matrix and Pearson correlation matrix.

### Identification of TAD boundary

Insulation score was calculated in each 40 kb bin by GENOVA tools, then local minimum search within 120 kb was performed. We assigned 40-kb bins with local minimum to TAD boundaries if the score differences between bins local minimum and bins with local maxima was greater than 0.3. In this condition, we got a high reproducibility of TAD boundary between biological replicates (>88%).

### ChIP-seq analysis

Adaptor sequences and low-quality bases in reads were trimmed using Trim Galore version 0.3.7 (http://www.bioinformatics.babraham.ac.uk/projects/trim_galore/). Then trimmed reads were aligned to the mouse GRCm38 genome assembly using bowtie version 0.12.7 (Langmead, Trapnell, Pop, & Salzberg, 2009) with default parameters. Duplicated reads were removed using samtools version 0.1.18 (Li et al., 2009).

### RNA-seq analysis

Adaptor sequences and low quality bases in reads were trimmed using Trim Galore version 0.3.7 (http://www.bioinformatics.babraham.ac.uk/projects/trim_galore/). To calculate RPM values in each 250 kb bin, trimmed reads were aligned to the mm10 genome build using bowtie version 0.12.7 with –m 1.

Duplicated reads were removed using samtools version 0.1.18. The number of mapped reads in each 250 kb bin was counted by featureCounts (Liao, Smyth, & Shi, 2014), then, RPM values in each 250 kb bin was calculated. To identify differentially expressed genes or repeats, the trimmed reads were mapped to the mouse GRCm38 genome assembly using TopHat (v2.1.1) with –g 1 (Trapnell, Pachter, & Salzberg, 2009). After read mapping, the number of reads mapped in genes or repeats was counted by TEtranscripts (v1.4.11) with default parameters) (Jin, Tam, Paniagua, & Hammell, 2015). We performed two biological replicates for RNA-seq and identified DE genes and repeats by DESeq2 (adj. P-value < 0.05, FC > 2) (Love, Huber, & Anders, 2014).

### Visualization of NGS data

The Integrative Genomics Viewer (IGV) (Robinson et al., 2011) was used to visualize NGS data. For Hi-C contact matrix and correlation matrix, we used Juicer Tools (Durand et al., 2016).

### Statistical analysis

All methods for statistical analysis and P-values in this study are described in each figure legend and included in each figure, respectively.

## Data availability

The NGS data used in this study is listed in Supplementary Table 1.

## ACKNOWLEDGEMENTS

We thank the staff of the Support Unit for Bio-Material Analysis (BMA) at the RIKEN Center for Brain Science (CBS) Research Resources Division (RRD) for NGS library construction, DNA sequencing and flow cytometry. We would also like to thank our colleagues at Shinkai laboratory for their support and valuable comments.

## Author contributions

K.F. and Y.S. designed and conceived the study. K.F. and Y.S. supervised the study and interpreted the data. C.S. performed molecular and cellular experiments and generated the ChIP-seq, RNA-seq and Hi-C-seq libraries. K.F. performed informatics analysis of generated NGS data. K.F. and Y.S. wrote the manuscript and prepared figures. All authors read, discussed, and approved the manuscript.

---

**Supplementary Figure S1.**
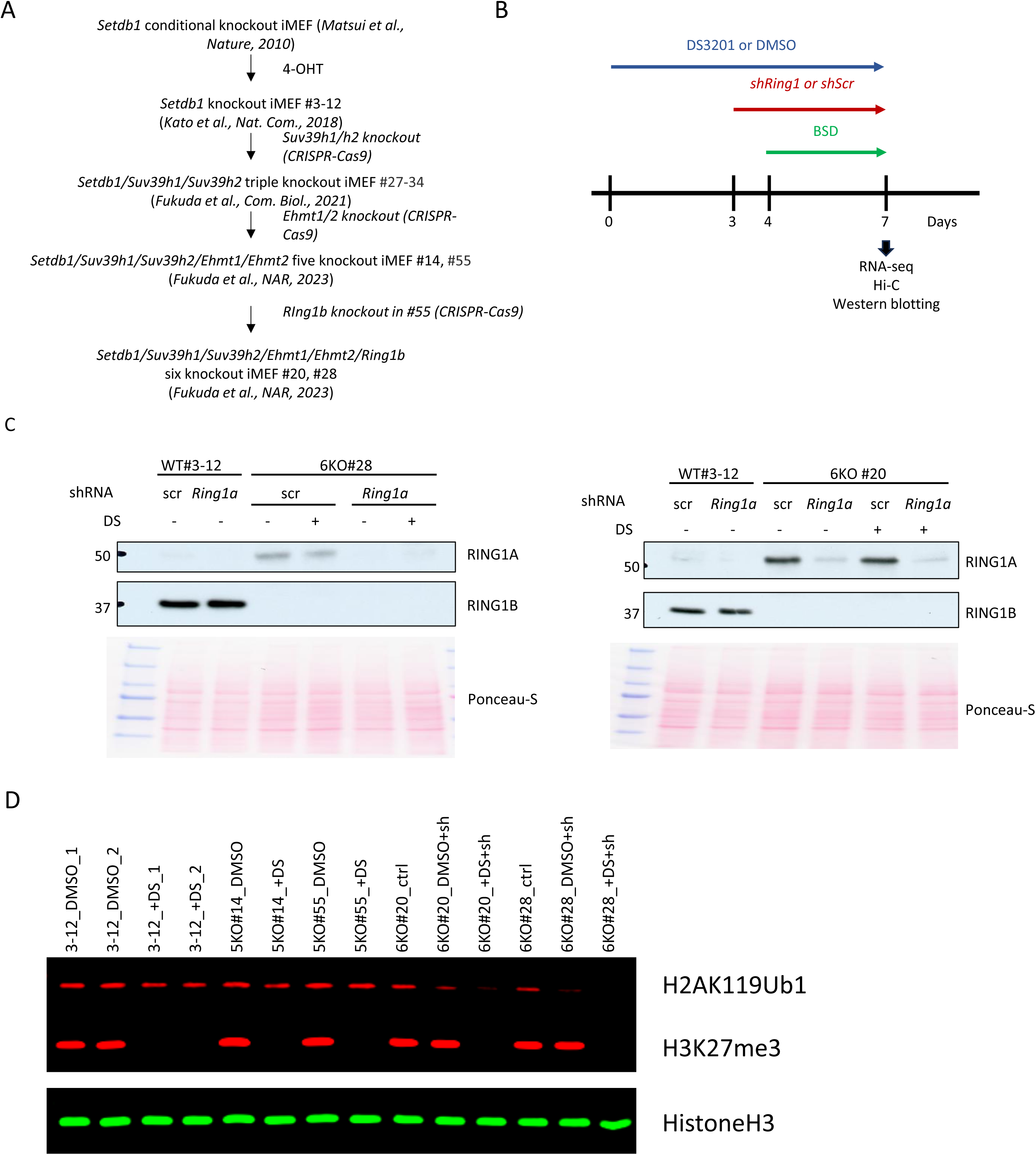
Establishment of H3K9 methylation/H3K27me3/uH2A depleted iMEFs. (A) Flow chart illustrating the process of generating 6KO iMEFs. We established two 6KO clones (#20, #28) from 5KO iMEFs. (B) Experimental design of establishment of H3K9 methylation/H3K27me3/uH2A-depleted iMEFs. (C) Western blotting analysis of RING1A and RING1B. *Ring1a*-shRNA treatment efficiently decreased RING1A expression in 6KO iMEFs. (D) Western blotting analysis of uH2A and H3K27me3. Both DS3201 and *Ring1a-*shRNA treatment efficiently decreased uH2A and H3K27me3 in 6KO iMEFs.

**Supplementary Figure S2.**
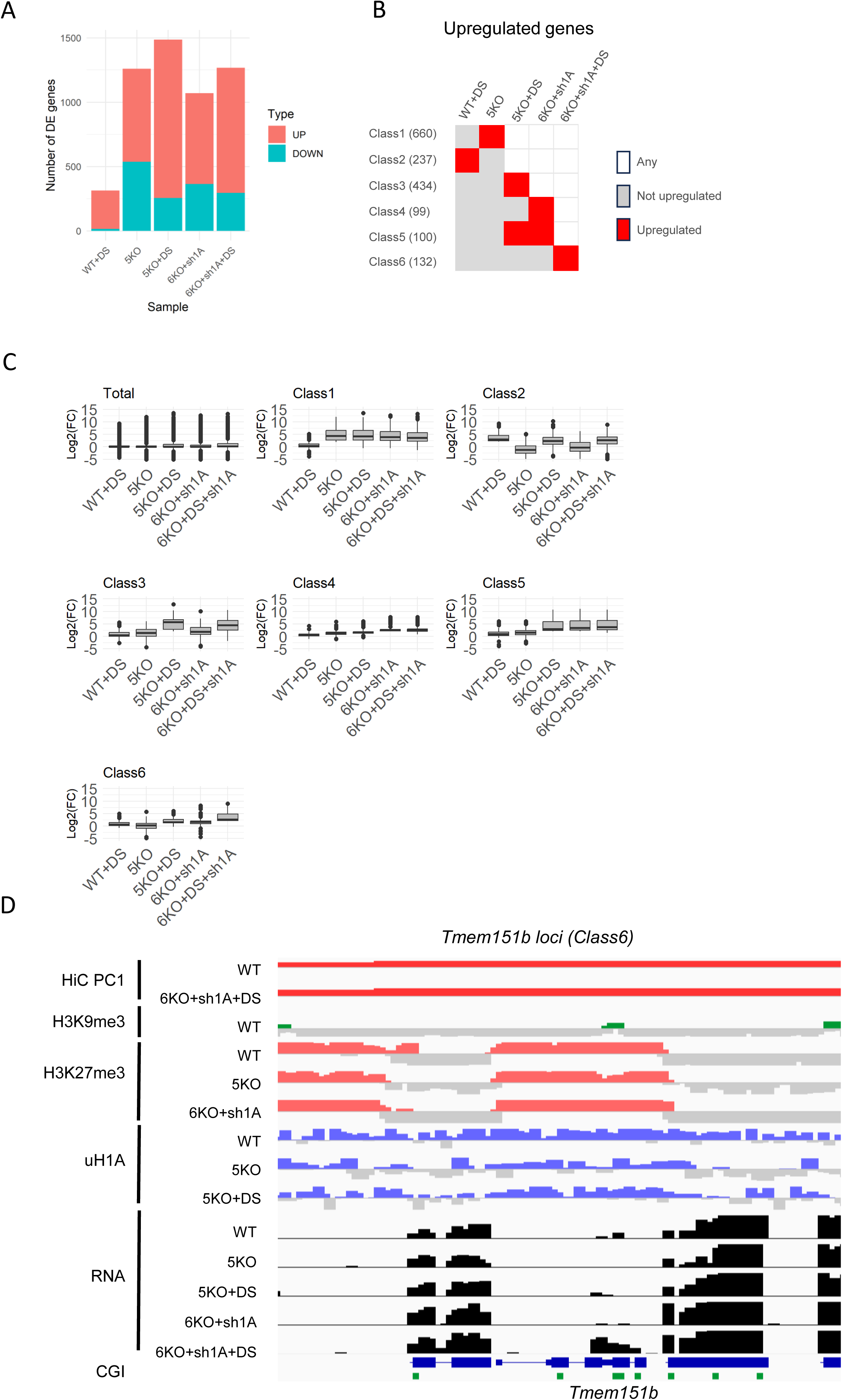
Regulation of gene repression by the H3K27me3 and RING1A/B after the loss of H3K9 methylation. (A) Bar plots showing the number of DE genes in each sample. (B) Classification of upregulated genes based on different combinations of upregulated samples. Red, grey, and white represent upregulated, non-upregulated, and any status, respectively. (C) Boxplots showing expression change of upregulated genes. (D) Representative region of Class 6 gene. uH2A and H3K27me3 are redistributed to *Tmem151b* loci in 5KO iMEFs and are independently maintained each other. uH2A and H3K27me3 repress *Tmem151b* without compartment change.

**Supplementary Figure S3.**
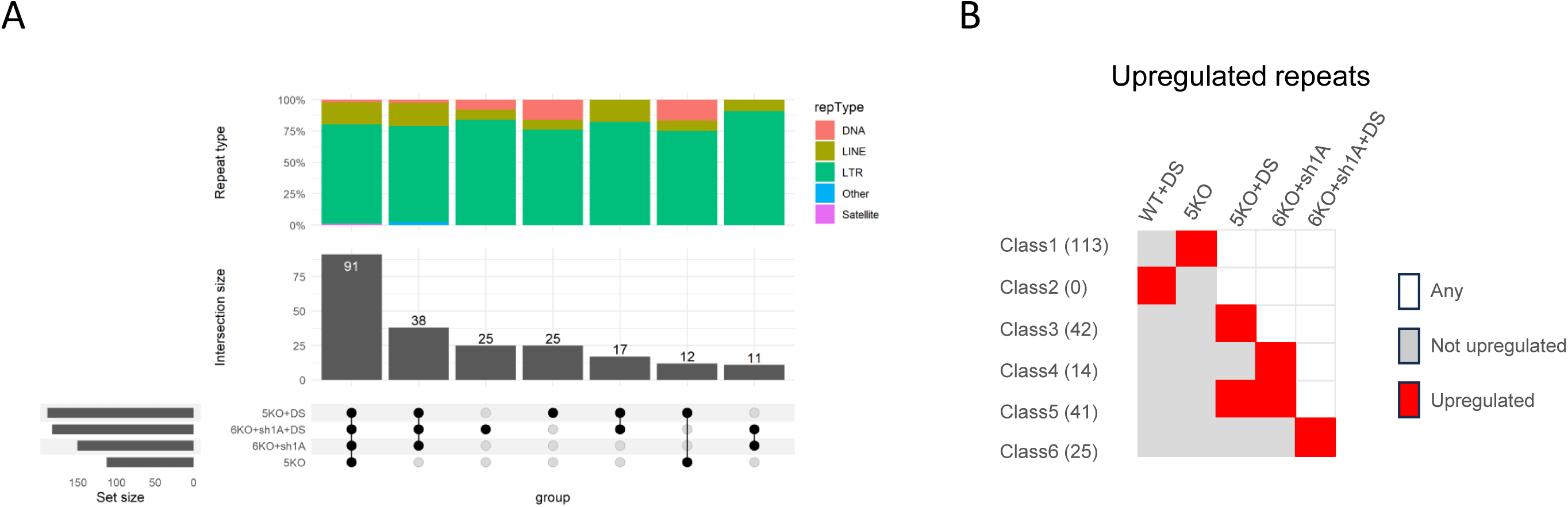
Regulation of transposon repression by the H3K27me3 and RING1A/B after the loss of H3K9 methylation. (A) Upset plot of upregulated transposons. Bar plots show the fraction of repeat type of each group. (B) Classification of upregulated genes based on different combinations of upregulated samples. Red, grey, and white represent upregulated, non-upregulated, and any status, respectively.

**Supplementary Figure S4.**
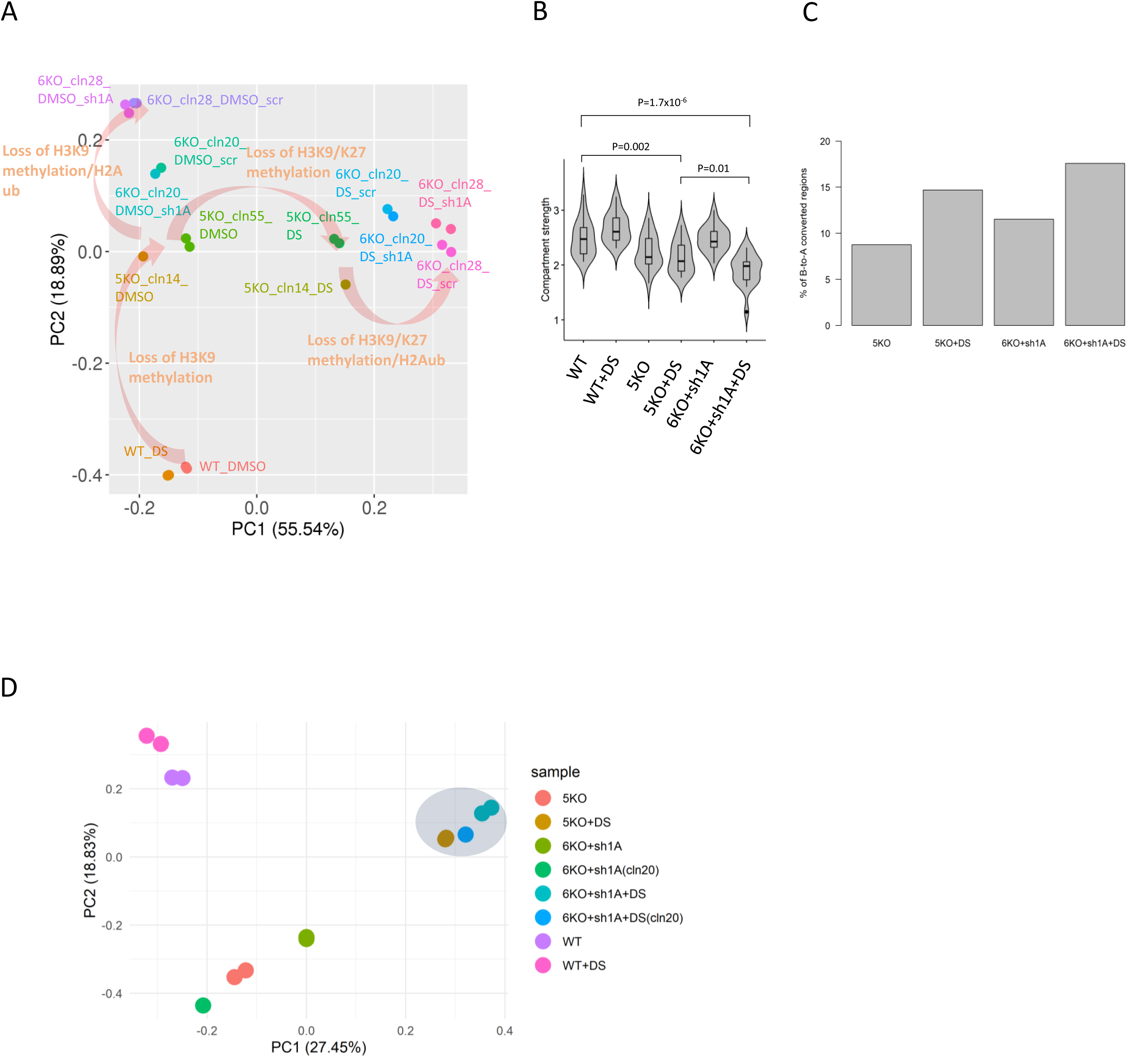
H3K27me3 and RING1A/B maintain A/B compartment segregation after the loss of H3K9 methylation. (A) PCA plot of Hi-C PC1 values of 250-kb bin. (B) Violin plots of compartment strength. P-values were calculated by Tukey’s test. (C) Fraction of B-to-A converted 250-kb bin. (D) PCA plot of insulation score of 40-kb bin.

